# Expression of a RAS degrader via synthetic nanocarrier-mediated mRNA delivery reduces pancreatic tumors

**DOI:** 10.1101/2024.06.11.598439

**Authors:** Taylor E. Escher, Simseok A. Yuk, Yuan Qian, Wenan Qiang, Sultan Almunif, Swagat Sharma, Evan A. Scott, Karla J. F. Satchell

**Author notes:** These authors contributed equally (Taylor E. Escher, Simseok A. Yuk). T.E.E., University of Kansas, Lawrence, KS, USA, S.A.Y., Merck & Co., West Point, PA, USA, Y.Q., Innocare Pharma, Beijing, China, E.A.S, University of Virginia, Charlottesville, NC, USA. Karla J. F. Satchell: Northwestern University, Feinberg School of Medicine, Chicago, Illinois, 60611, United States. Evan A. Scott: University of Virginia, School of Medicine and School of Engineering and Applied Science, Charlottesville, North Carolina, 22904.

## Abstract

Therapeutic gene expression can address many of the challenges associated with the controlled delivery of intracellularly active biologics, such as enzymes that degrade RAS for treatment of RAS-driven cancers. Here, we demonstrate that an optimized synthetic nonviral gene delivery platform composed of poly(ethylene glycol)-*b*-poly(propylene sulfide) (PEG-PPS) can block copolymers conjugated to a dendritic cationic peptide (PPDP2) for nontoxic delivery and therapeutic expression of mRNA within human pancreatic cancer cells and tumors. The naturally occurring bacterial enzyme RAS/RAP1-specific endopeptidase (RRSP) is a potent RAS degrader that specifically targets all RAS isoforms. Using PPDP2, *rrsp*-mRNA is delivered to human pancreatic cells resulting in RRSP protein expression, degradation of RAS, and loss of cell proliferation. Further, pancreatic tumors are reduced with residual tumors lacking detectable RAS and phosphorylated ERK. Using structural modeling, we further demonstrate that a noncatalytic RAS-binding domain of RRSP provides high specificity for RAS. These data support that the synthetic nanocarrier PPDP2 can deliver *rrsp*-mRNA to pancreatic tumor cells to interrupt the RAS signaling system.

## INTRODUCTION

RAS proteins control cell proliferation in both normal and cancerous cells. Currently, the Federal Drug Administration approved RAS-directed therapies specifically target Kirsten rat sarcoma (KRAS) mutants, which are present in about 30% of all cancers. These molecules in clinical use against lung and other cancers have demonstrated a high propensity for driving drug resistance, leading researchers to consider alternative strategies for RAS-targeted therapies.^1^ Cutting-edge strategies to target RAS driven cancers include “RAS Degraders”, which specifically target RAS for proteolytic turnover and result in lowered levels of RAS within cancer cells that can be useful in treating nearly all tumors.^2, 3^ In addition, “pan-RAS” degraders target all forms of RAS in the cell, devoiding cells of RAS and thereby stopping cell proliferation.^2, 3^ While the vast majority of RAS degraders are small molecules, intracellularly active enzymes for direct RAS proteolysis are an underexplored burgeoning category that is limited by challenges with efficient and non-immunogenic controlled delivery of biologics into cancer cells.^3^

The RAS/RAP1-specific endopeptidase (RRSP) is a well-studied intracellular RAS degrader. Originally termed DUF5, RRSP is a bacterial cytotoxic effector domain from the multifunctional-autoprocessing repeats-in-toxin (MARTX) toxin.^4^ RRSP site-specifically cleaves RAS and its close homologue RAP1, between residues tyrosine-32 and aspartic acid-33 within the Switch I region, thereby preventing interaction with RAF kinases in the RAS-ERK signaling axis. RRSP has been shown to be highly specific and does not target other closely related GTPases.^5, 6^ RRSP can cleave all three of the major RAS isoforms (H, N, and K), both GTP and GDP-bound RAS, as well as the most common oncogenic RAS mutations, including G12C, G12D, G12V, G13D, and Q61R.^5–7^ Within cells, RRSP degradation of RAS leads to G1 cell cycle arrest that can progress to apoptosis, senescence, and loss of cell proliferation in more than 80% of all cell lines where it has been tested, including leukemia, non-small cell lung carcinoma, colorectal carcinoma, central nervous system cancers, melanoma, ovarian cancers, renal cancers, pancreatic cancer, and breast cancer.^4,7–9^ The major limitations with the use of RRSP as a cancer therapeutic are its 56 kilodalton size and that the active domain cannot transit across the cell plasma membrane without the remaining portions of the larger toxin. The advent of nucleic acid delivery to cells provides a potential strategy for RRSP to be expressed within cells for RAS degradation and loss of cell proliferation.^10, 11^

Efficient and scalable methods have been described for loading bioactive molecules, including both proteins and nucleic acids, within synthetic nanocarriers composed of the self-assembling polymer poly(ethylene glycol)-*b*-poly(propylene sulfide) (PEG-PPS).^12–14^ PEG-PPS synthetic nanocarriers have been tested across diverse disease models,^10, 15, 16^ and validated as non-immunogenic in human blood.^17^ The platform has been demonstrated to be noninflammatory and nontoxic in nonhuman primates^18^ and humanized mice.^14^ We recently engineered a PEG-PPS copolymer variant for stable nontoxic delivery of nucleic acids by linking a cationic dendritic peptide (DP) tertiary block via a reduceable bond to generate PPDP (Figure 1A).^11^ After an optimization that tested nearly a dozen PPDP variants, the “PPDP2” derivative was found to complex with plasmid DNA and assemble into highly stable ∼100 nm vesicular nanocarriers.^11^ Furthermore, PPDP2 undergoes pH-dependent disorder-to-order transitions to adopt a unique helical conformation under acidic conditions that promotes both the endosomal escape and efficient release of plasmids to the cytoplasm.^11^ This optimized nanocarrier had exceptionally low toxicity compared to alternative nonviral synthetic platforms, such as polyethyleneimine and transfects cells under standard in vitro culture conditions in the presence of serum.^11^ Building upon these prior in vitro studies for plasmid delivery, here, we demonstrate that PPDP2 can also deliver mRNA for in vivo expression of a biologic within nonphagocytic pancreatic cells and tumors. Further, when an RRSP encoding *rrsp*-mRNA is delivered via PPDP2, the RRSP enzyme is highly expressed, cleaves RAS, and leads to loss of cell proliferation and tumor regression.

**Figure 1.**
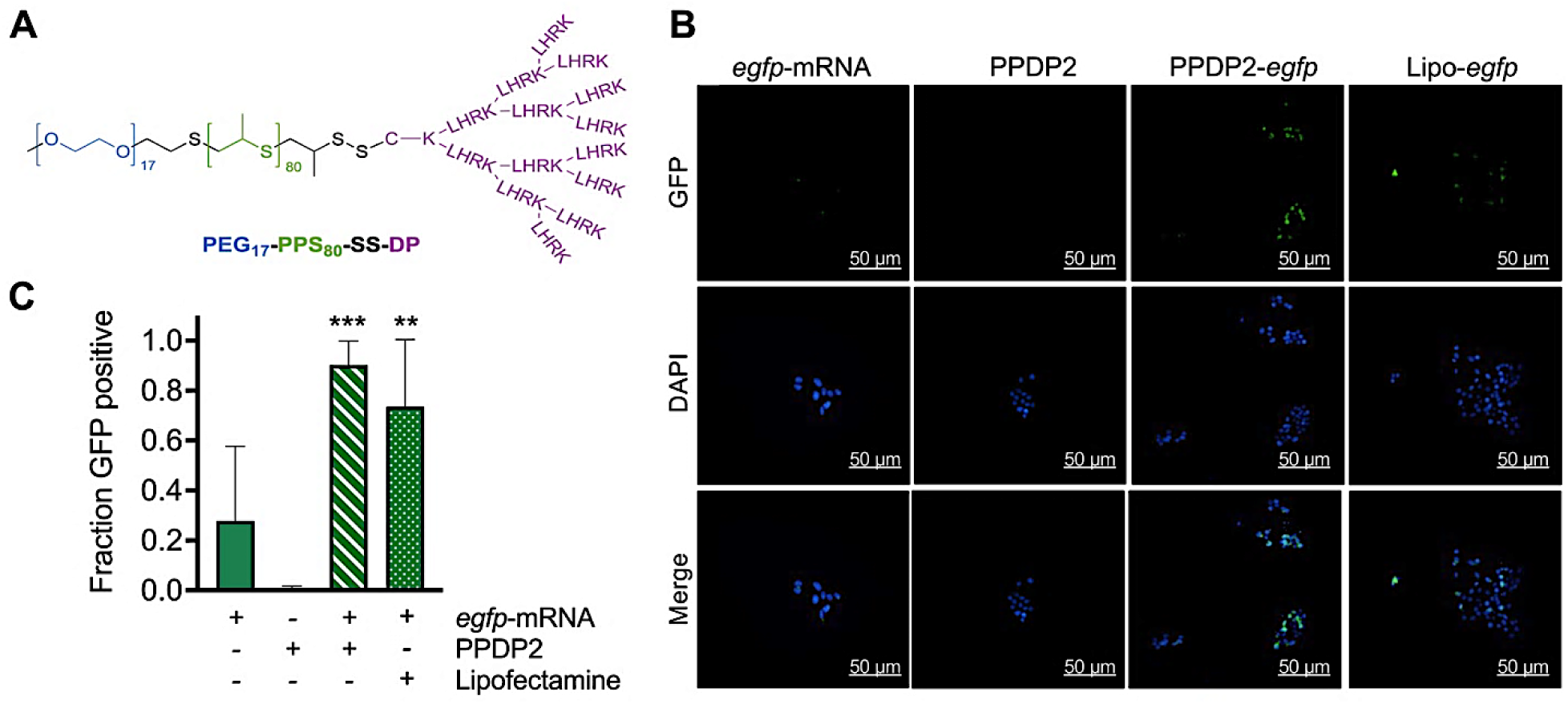
PPDP2 nanocarriers deliver *egfp*-mRNA into pancreatic cancer cells. (A) Schematic of PPDP2 nanocarrier chemistry. (B) Representative fluorescent images of PANC-1 cells after transfection of cells with 1 µg of *egfp*-mRNA alone, PPDP2 alone (1:40 w/v%) (PPDP2), *egfp*-mRNA (1 µg) with PPDP2 (PPDP2-*egfp*) or *egfp*-mRNA (1 µg) with MessengerMAX Lipofectamine (Lipo-*egfp*). (C) Quantification of the fraction of EGFP positive cells (green) from five imaged frames are shown as a histogram. P value calculated using a one-way ANOVA and Dunnett’s multiple comparisons test assuming normal distribution, \*\**p*<0.01, \*\*\**p*<0.001.

## RESULTS

### Nanocarrier delivery of mRNA to non-phagocytic cells

Our prior investigation of nucleic acid loading and delivery via the optimized PEG-PPS copolymer PPDP2 focused on delivery of plasmid DNA to phagocytic cells in vitro.^11^ Thus, our first objective for this study was to demonstrate PPDP2 nanocarrier delivery of mRNA into nonphagocytic cancer cells in culture. In human pancreatic PANC-1 cells, PPDP2 nanocarriers delivered *egfp-*mRNA (1 µg) to cancer cells resulting in expression of eukaryotic-optimized green fluorescent protein (EGFP) (Figure 1B). The number of EGFP-positive cells showed high efficiency, with ∼80% of cells transfected. The efficiency of transfection was 2.5-fold higher than cells treated with *egfp*-mRNA alone and was similar to transfection of *egfp*-mRNA using MessengerMAX lipofectamine (Figure 1C).

### Delivery of *rrsp*-mRNA for expression of RRSP RAS degrader

Next, we sought to determine whether we could employ PPDP2 synthetic nanocarriers for delivery of *rrsp*-mRNA for expression of the RRSP RAS degrader within cells. Human pancreatic PANC-1 cells were treated with varying concentrations of *rrsp*-mRNA using either PPDP2 or lipofectamine as the carrier. Treated cells showed significant concentration-dependent loss of cell proliferation, resulting in low cell concentration by 24 hours following transfection.

At concentrations as low as 1.25 µg of *rrsp-*mRNA delivered by PPDP2, there was greater than 60% reduction in cells as measured by crystal violet staining (Figure 2A). To demonstrate impact on RAS, we detected pan-RAS levels following treatment for only 3 hours followed by 21 hours of incubation to allow for protein expression while preserving cell viability. We observed reduced RAS levels in cells treated with the *rrsp*-mRNA, even when values were adjusted for low cell recovery (Figure 2B). Note, PPDP2 in the absence of mRNA led to a variable response with either a slight increase or decrease in RAS levels across different experiments.

**Figure 2.**
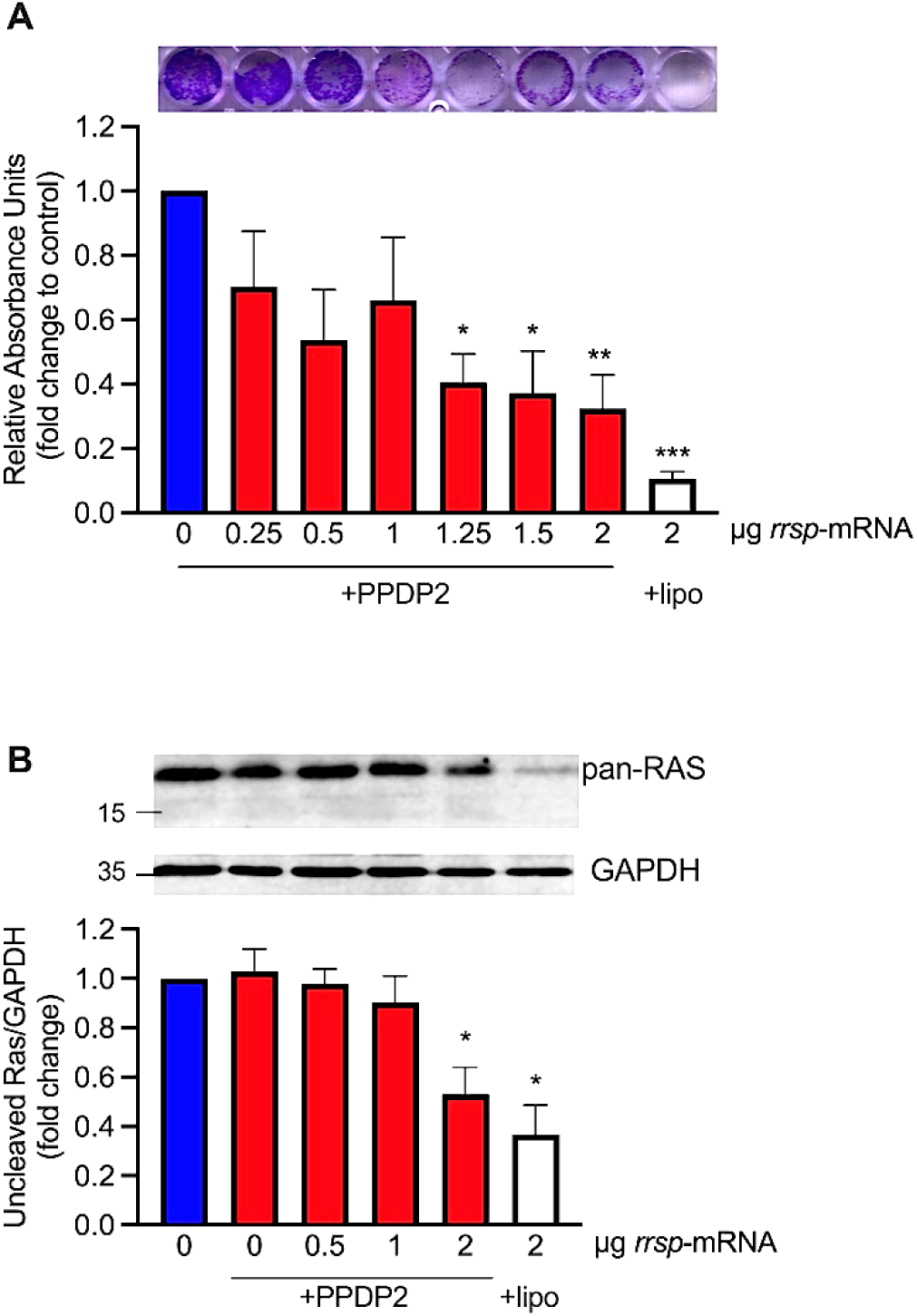
PPDP2-*rrsp*-mRNA impacts cell proliferation and reduces RAS levels in human pancreatic cancer cells. (A) Representative crystal violet staining and spectrophotometer quantification (*n*=3) from PANC-1 cells treated as indicated. (B) Western blot and quantified expression levels of RAS using a pan-RAS antibody in cell lysates collected after transfection for 3 hours as indicated and followed by 21 hours incubation after media exchange. Densitometry quantification from replicate Western blots for RAS normalized to GAPDH control (*n*=4-5). P values were calculated using a one-way ANOVA and Dunnett’s multiple comparisons test assuming normal distribution, \**p*<0.05, \*\**p*<0.01, ***p<0.001

### Pancreatic tumor reduction following delivery of mRNA for expression of RRSP

To test for delivery of mRNA in the context of a tumor in vivo, *mCherry*-mRNA was delivered by PPDP2 into PANC-1 xenograft tumors. PANC-1 pancreatic cell line-derived xenograft tumors were first established in immunodeficient *nu/nu* mice, then treated by intratumoral (i.t.) injection with PPDP2/*mCherry-*mRNA. After four weeks of treatment three times per week (excluding weekends), resected tumors showed high levels of expression of mCherry by Western blotting in 2 out of 3 mice (Figure 3A) and by immunohistochemistry (IHC) staining (Figure 3B) showing 20-60% of cells positive for mCherry expression.

**Figure 3.**
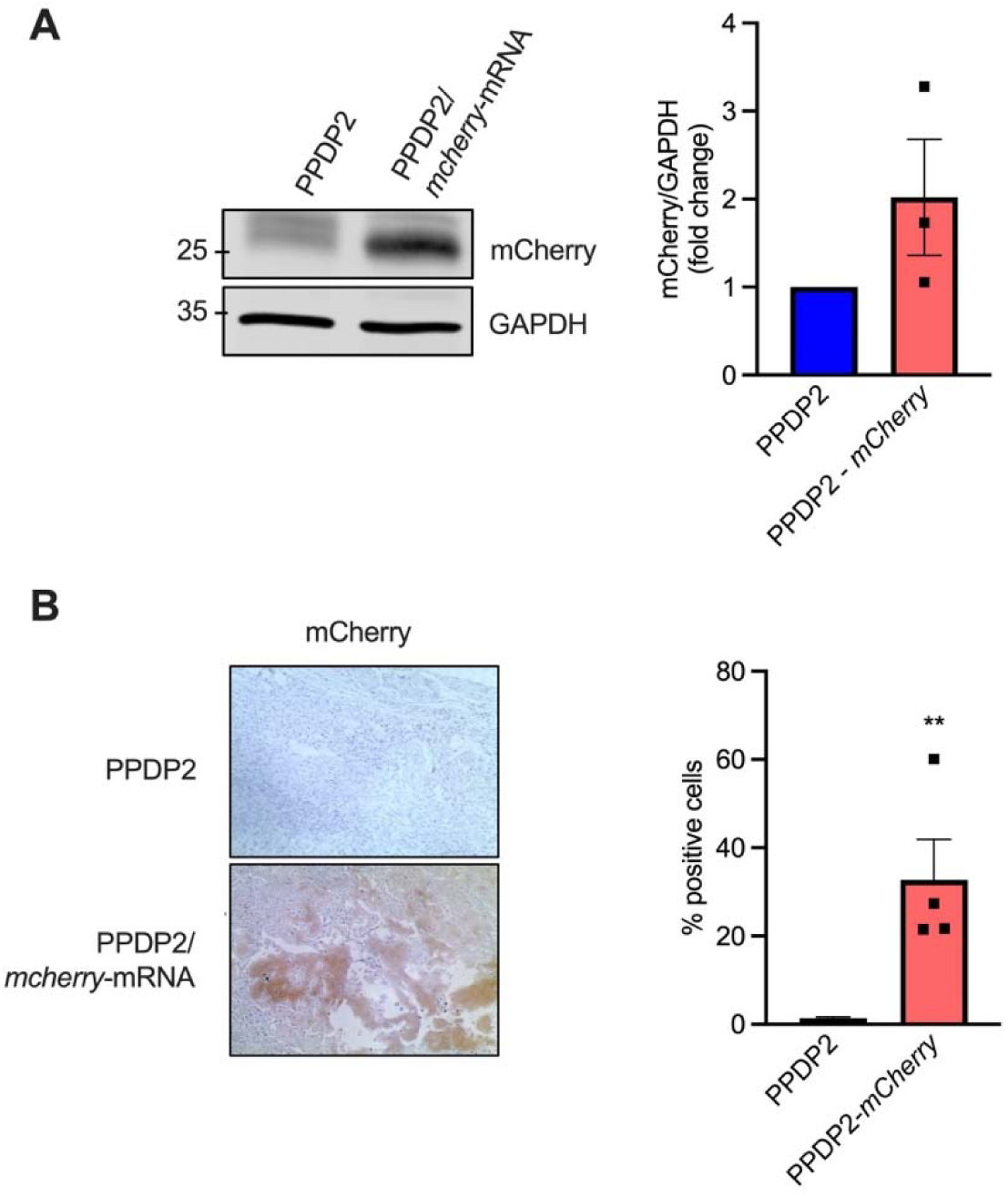
PPDP2 nanocarriers deliver *mCherry-*mRNA into PANC-1 xenograft tumors. (A) Representative Western blot and quantification of mCherry levels from PANC-1 xenografts injected with PPDP2 alone (PPDP2) or PPDP2–*mCherry*-mRNA (PPDP2-*mCherry*) (*n*=3). (B) mCherry IHC staining and quantification from PANC-1 xenografts injected as indicated (*n*=4). P values were calculated using Student’s *t* test, \*\**p*<0.01.

We next asked whether PPDP2-*rrsp-*mRNA affects pancreatic duoductal adenocarcinoma (PDAC) tumor xenograft growth in mice. For these studies, the *rrsp*-mRNA is identical to that used for cell line-derived xenograft studies except for the addition of codons for a hemagglutinin (HA) tag for antibody detection of the protein expressed in tumors. In addition, an *rrsp*\*-mRNA that expresses the H451A mutant, and thus cannot cleave RAS, was added as an additional control. PANC-1 xenograft tumors were established and then treated by intratumoral (i.t.) injection in two independent experiments. In experiment 1, xenograft tumors were treated three times per week with 11 total injections over 27 days (Figure 4A). In experiment 2, xenograft tumors were treated every other day for 11 total injections over 22 days (Figure 4B). Both experiments resulted in a significant difference in the growth rate of tumors in the PPDP2-*rrsp*-mRNA treatment group compared to the PPDP2 group alone (Figure 4A,B). At day 22 for both experiments, 8/10 mice in the treatment group showed tumor regression with one tumor fully resolved (Figure 4D,E).

**Figure 4.**
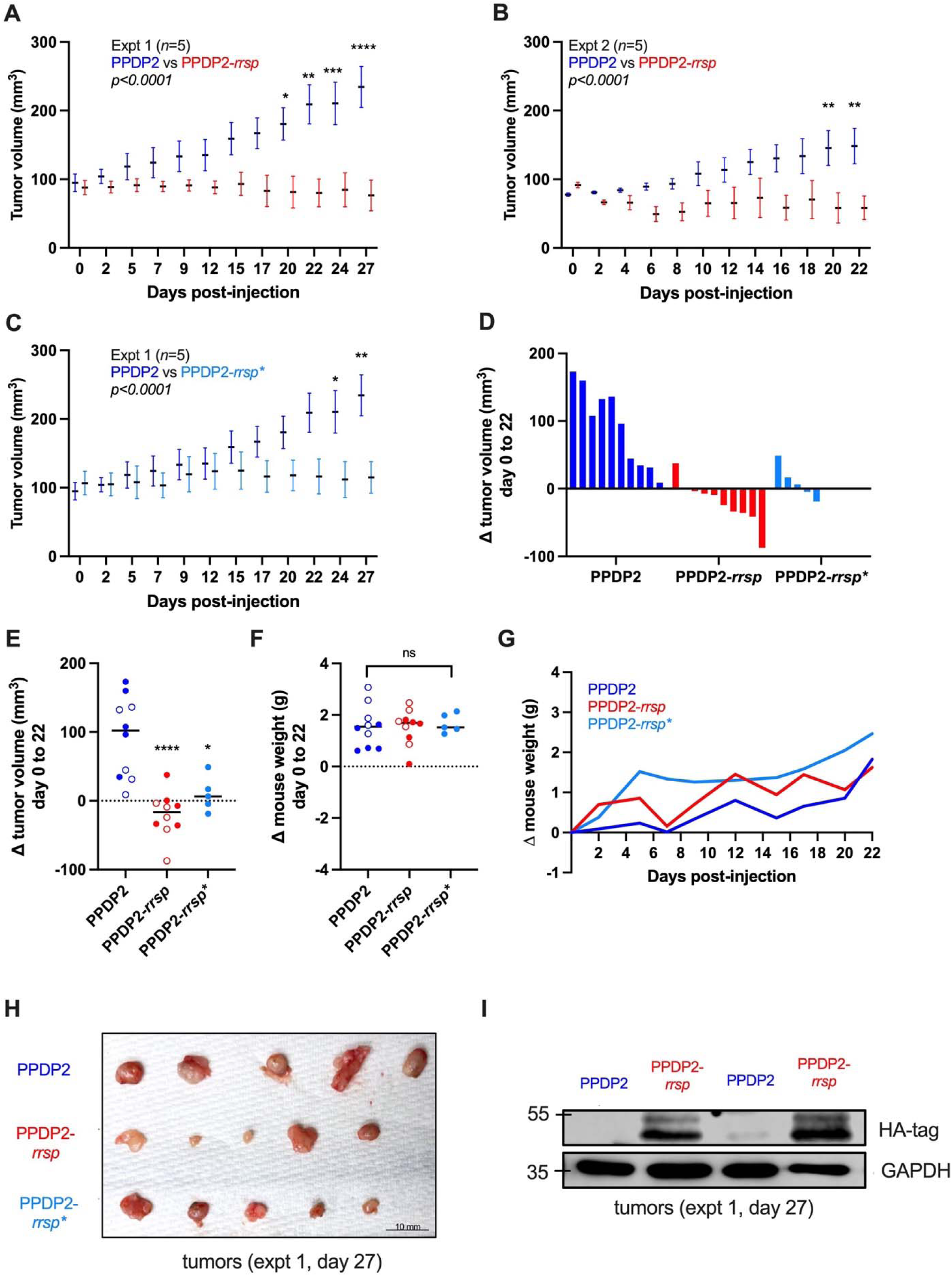
PPDP2-*rrsp*-mRNA induces regression of PANC-1 xenograft tumors. (A-C) Mean +/-SEM of tumor volume measured over time as indicated with day 0 representing the tumor size just before the first treatment. Mice were newly injected after each measurement was taken. PPDP2 control in panel C is the same as panel A as experiments were conducted at the same time but separated for clarity of presentation. Statistical analysis conducted using two-way ANOVA shows significant difference in growth rate between groups over time (*p*<0.0001). (D,E) Change in size of individual tumors relative to Day 0 following, PPDP2, PPDP2/*rrsp*-mRNA, and PPDP2/*rrsp*\*-mRNA treatment were calculated and presented as waterfall plot for evaluation of tumor disease progression (D) and scatter plot for statistical comparison (E). All tumors from day 0 to day 22 with closed circles representing Experiment #1 and open circles representing Experiment #2. (F,G) Weight of individual mice at day 22 (F) and average weight over time (G). Individual plots for each mouse over time in Supporting Figure 1. (H) Photograph of resected tumors from Experiment 1, day 27. (I) Western blot of RRSP-HA in two tumors selected from Experiment 1. Remaining 3 tumors were used for IHC shown in Figure 5. Statistical difference of treatment group compared to PPDP2 over time in panels A-C determined by Two-Way RM ANOVA with Sidak’s multiple comparison test. Comparison of treatment group to PPDP2 measurement for Panels E,F by Student’s *t* test. Significance indicated by asterisks: \**p*<0.05, \*\**p*<0.01, \*\*\**p*<0.001, \*\*\*\**p*<0.0001

There was no significant change in mouse weight at day 22 (Figure 4F) or over the course of the experiment (Figure 4G and Supporting Figure 1) indicating no generalized toxicity. Four out of five tumors in the treatment group resected from mice in experiment 1 were visibly smaller at day 27 (Figure 4H). For experiment 2, mice were extended without treatment for additional 23 days. Mice in the control group showed continued slow growth while mice in the treatment group showed 2 continued to regress while 2 tumors showed re-growth indicating variable response to the treatment (Supporting Figure 2A,B).

Two representative tumors treated with PPDP2-*rrsp*-mRNA showed high expression of RRSP in tumor tissue by western blotting (Figure 4I). The management of tumor growth by delivery of *rrsp*\*-mRNA was surprising (Figure 4C), suggesting that the expressed level of RRSP* over time accumulates in tumor cells to a level sufficiently high to bind RAS and inhibit its activity without enzymatic cleavage.

Xenograft tumor tissue analyzed by histology, trichrome, and immunohistochemical staining, and showed control PANC-1 tumors treated with only PPDP2 had enlarged nuclei and cytoplasm with dense collagen layers (Figure 5A), along with a high expression level of the CK-19 cytokeratin marker indicative of PDAC (Figure 5B) and of the Ki-67 proliferation marker (Figure 5C). Both RAS and phospho-ERK levels were high, consistent with active tumor growth (Figure 5D,E). In contrast, tumors treated with PPDP2-*rrsp*-mRNA showed reduced cell cytoplasm, immune cell infiltration, and loss of tumor tissue density or organization (Figure 5A). CK-19 and Ki-67 levels were reduced or absent (Figure 4B,C). Most notably, RAS and phospho-ERK levels were significantly reduced, which is indicative of target engagement (Figure 5D,E). Notably the PPDP2-*rrsp*\*-mRNA group did not show reduced RAS levels (Figure 5D), which is consistent with the expressed RRSP* protein not being able to cleave RAS. There was however a statistically significant loss of phospho-ERK in tumor tissue following treatment with *rrsp*-* mRNA (Figure 5E), consistent with the suggestion that catalytically inactive RRSP* may be binding to RAS and inhibiting its downstream effect on ERK. Overall, our data show that both *rrsp-*mRNA and *rrsp*-*mRNA delivered through PPDP2 nanocarriers significantly inhibit RAS signaling, likely by different mechanisms, with both resulting in tumor regression.

**Figure 5.**
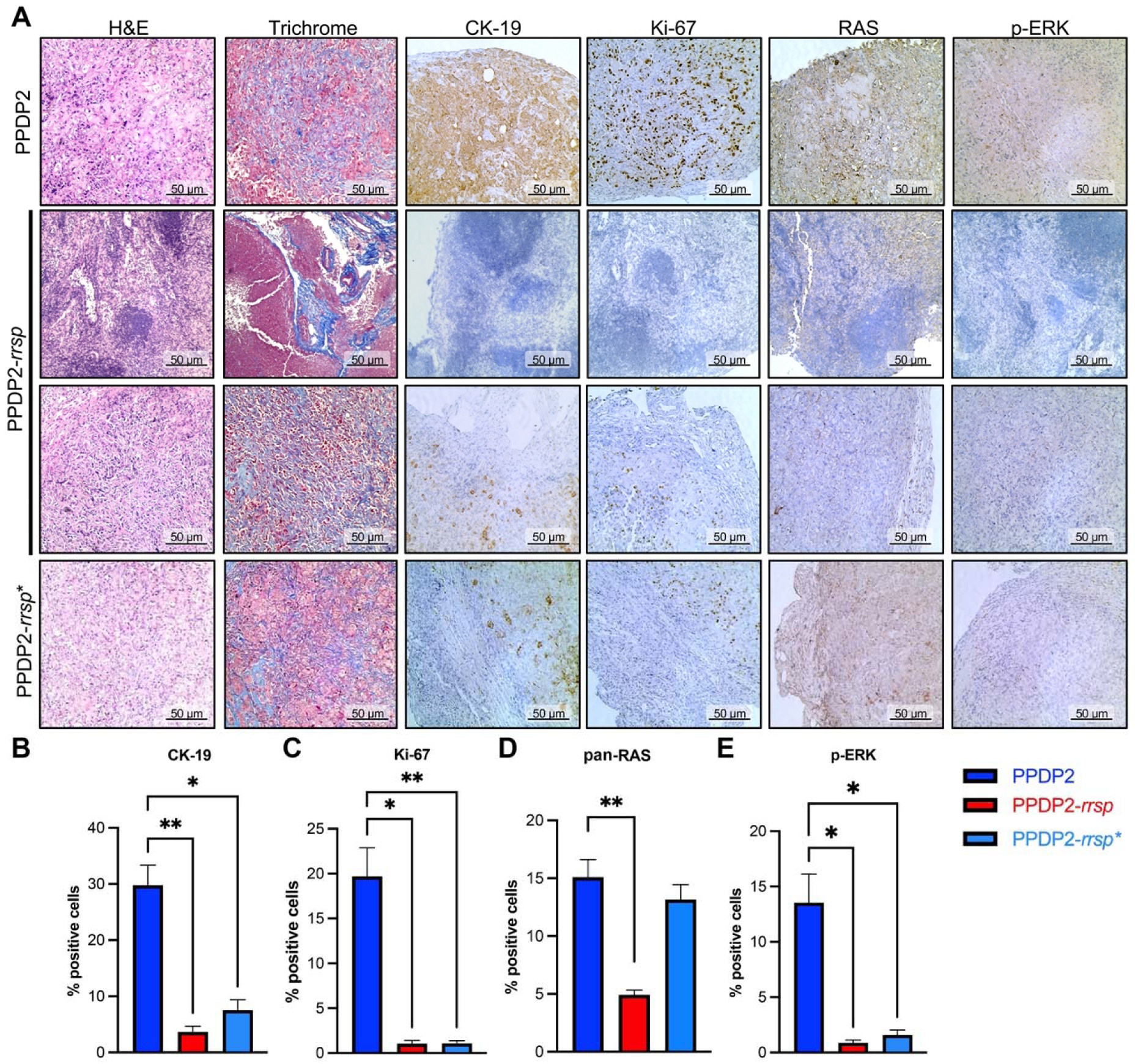
PPDP2*-rrsp-*mRNA impact on tissue organization and protein expression. (A) H&E, Masson’s trichrome, and IHC staining with anti-CK-19, anti-Ki-67, anti-pan-RAS, and anti-phospho-p44/42 MAPK. (B-E) ImageJ was used to quantify intensity of brown staining as indicated at top (*n=*5). Data were presented as mean ± SEM. *P* values were calculated using a one-way ANOVA and Dunnett’s multiple comparisons test, assuming normal distribution * *p* < 0.05, ** *p* < 0.01.

## DISCUSSION

Overall, our data demonstrate that PPDP2 can deliver mRNA to nonphagocytic cancer cells and tumors, resulting in expression of fluorescent proteins EGFP and mCherry. Further, delivery of *rrsp-*mRNA via PPDP2 achieved cytotoxicity and loss of proliferation and reduced RAS levels in surviving cells. Growth of pancreatic tumors was also inhibited, and regression occurred in 8/10 tumors tested. PPDP2 demonstrated a remarkable ability to achieve these results with no detectable toxicity despite four weeks of administration every other day, which was demonstrated by no weight loss or signs of distress in the animals.

An unexpected result suggests the in vivo study was likely conducted with more PPDP2 and/or mRNA than necessary, as evidenced by the high expression levels of mCherry in some animals and also the RRSP proteins. Indeed, the expression of inactive RRSP* was sufficient to inhibit tumor growth in most mice. To understand why RRSP could impact RAS function without cleaving, we generated an AlphaFold2 (AF2) model of the dimeric structure of RRSP bound to KRAS (Figure 6A). RRSP is comprised of a membrane targeting C1 domain and the large C2 domain formed as two lobes (termed C2A and C2B). The AF2 model predicts that the C2 domain binds the RAS G-domain with RAS residues 21-35 comprising the KRAS Switch 1 inserted into the active site (Figure 6A). Unexpectedly, both the RRSP C2A and C2B domains contact KRAS. The C2B catalytic domain contacts Switch 1 and Switch 2 as expected since Switch 1 is cleaved by RRSP. However, the model further predicts α8, α9, and α10 helices of the noncatalytic RRSP C2A domain (Figure 6B) contacts the RAS G-domain β-sheet comprised of β1-β2-β3 and α2 (Figure 6C) revealing a major binding face of the protein. The contacted residues within RAS are highly conserved across all RAS proteins known to be cleaved by RRSP. This dual contact AF2 model is supported by prior data that overexpression of C2A alone in cells is cytotoxic when overexpressed, whereas overexpression of C2B alone is not cytotoxic in the absence of C2A.^4^ Hence the high specificity of RRSP binding for RAS is dictated by the non-catalytic lobe of the protein, and thus our model supports that RRSP* could bind RAS and impede its activity, even as it cannot cleave the protein. Our data herein support that this binding is sufficient to reduce tumor growth.

**Figure 6.**
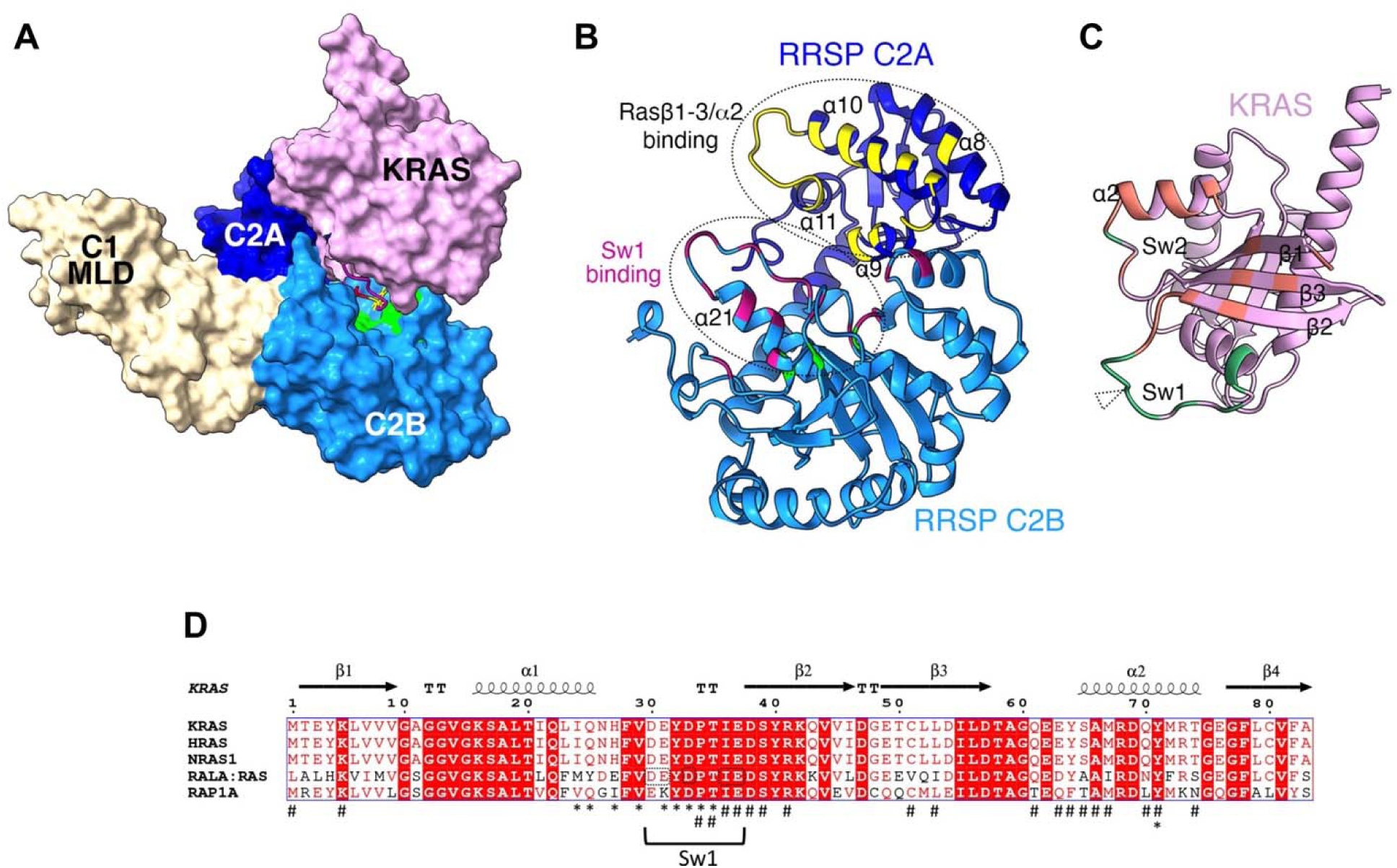
AF2 Structural model of RRSP in complex with KRAS suggests extensive contact sites. (A) Space filling AlphaFold2-generated model of RRSP (MARTX toxin aa 3594 – 4078) in complex with KRAS (aa 1-175). RRSP C1 membrane localization domain (MLD, white), C2A (dark blue), C2B (medium blue), and KRAS (pink). Switch 1 (Sw1) is shown only as a ribbon in magenta with scissile bond residues D32 and Y33 as sticks in yellow. Catalytic residues are colored lime green. Note that the Sw1 is pulled into the active site of RRSP. (B) Ribbon cartoon of RRSP (with C1 MLD removed) with backbone colored as in Panel A. Residues mapped as binding to KRAS are colored (residues that bind C2A are yellow and that bind C2B are magenta) (C) Residues in KRAS mapped as binding to C2A are colored orange and to C2B are colored green. Scissile bond is marked with triangle. (D) Alignment of all RAS sequences experimentally validated as successfully cleaved by RRSP. RALA:RAS is a chimera with 4 aa changes that alters the noncleaved RalA Sw1 to match the KRAS Sw1 (changed residues outlined) and is cleaved with equal efficiency as KRAS. 27 residues mapped as binding RRSP C2A are indicated with a hashtag (#) and to RRSP C2B as an asterisk (*). Sw1 and Sw2 are marked in panels C and D.

Recently, another group has tested delivery of *rrsp-*mRNA except using lipid nanoparticles (LNPs) optimized for activity in the higher reactive oxygen species tumor microenvironment.^19^ Our delivery system does not need this specific environment for payload release. Of note, our PPDP2 platform has potentially similar advantages with respect to scalability, safety, and versatility compared to LNPs. PEG-PPS nanocarriers have been extensively validated as stable delivery vehicles for targeting diverse therapeutics to specific cells and tissues in a wide range of preclinical animal disease models,^11, 12, 14, 20^ including in nonhuman primates.^19^

The nontoxic transfection combined with the stability of the PPS membrane and the flexibility to modify for advanced tumor targeting makes PPDP2 well suited for *in vivo* applications. Our work presented here verifies that the prior findings for plasmid delivery also hold true for mRNA payloads and for delivery to cancer cells, highlighting the versatility of the PPDP2 platform for nucleic acid delivery in general.

## MATERIALS AND METHODS

### Chemicals, Protein purification, and Cell lines

All chemicals were from Sigma-Aldrich unless otherwise specified. A CleanCap eGFP-mRNA mRNA for expression of EGFP and an *rrsp*-mRNA for expression of RRSP and an H451A catalytically inactive negative control mRNA (*rrsp*\*-mRNA) were synthesized by TriLink BioTechnologies. mRNA sequences were capped with 5’ AG head (CleanCap®), hemagglutinin (HA)-tagged (as indicated) and polyadenylated tail and 100% pseudouridine (Supporting Table 1).

Cell lines were obtained from the National Cancer Institute RAS Initiative or collaborators. Cell lines were confirmed free of *Mycoplasma* using VenorGeM Mycoplasma Classic Endpoint PCR assay and were also subjected to short tandem repeat analysis using the AmpFLSTR Identifiler PCR Amplification Kit to authenticate the cell lines, comparing the results with information located at https://web.expasy.org/cellosaurus/. Cells were cultured at 37°C and 5% CO2 atmosphere. PANC-1 cells were grown in Dulbecco’s Minimal Eagle’s Medium (DMEM, American Type Culture Collection formulation) with 10% Fetal Bovine Serum (FBS) and 1% penicillin/streptomycin (P/S).

### Synthesis and Characterization of PEG-*b*-PPS-ss-DP Polymer

To synthesize and load synthetic PEG-PPS nanocarriers with mRNA, block co-polymers PEG_17_-*b*-PPS_80_-pyridyl disulfide were synthesized and conjugated to the cationic DP via disulfide exchange as previously described.^11^ Briefly, good manufacturing practice grade synthesis of PEG_17_-*b*-PPS_80_-pyridyl disulfide was performed in collaboration with the contract research organization Sequens Group, while the final DP-end capping was completed in-house within a clean room. The complete chemical synthesis, including the structures of all monomers, initiators, and DP molecules, is provided in Supporting Figure 3. Commercial synthesis of DP was performed by Peptide 2.0, Inc. Additionally, comprehensive high pressure liquid chromatography (HPLC) and mass spectrometry HPLC and MS analyses were conducted by Peptide 2.0 to further characterize the DP. The theoretical molecular weight of the dendritic peptide was 6708.28 g/mol (Reference No. 167527-001). Its composition comprised 23.08% hydrophobic, 0.00% acidic, 69.23% basic, and 7.69% neutral amino acids. Additionally, the mass spectrum (M+H) showed a value of 6709.25 g/mol, and HPLC analysis at 220 nm using a C18 column with a linear gradient determined the peptide’s purity to be 99.56%. To calculate DP conjugation efficiency, the vacuum-dried PPDP2 conjugates were dispersed in molecular biology-grade water and dialyzed using Slide-A-Lyzer Dialysis Cassettes (20K MWCO, Thermo Fisher Scientific) for 24 hours to remove unreacted peptide 11. The dialysate was then collected, lyophilized, and weighed, resulting in 80 ± 15 mg from an initial 240 mg, corresponding to approximately 33% DP conjugation efficiency. The PEG-*b*-PPS precursors and PPDP2 polymer were characterized by using proton nuclear magnetic resonance (^1^H-NMR) and gel permeation chromatography (GPC). ^1^H-NMR area under the curve (AUC) measurements were normalized to the three hydrogens within PEG-OCH_3_ to confirm 17 repeating units of PEG and 80 repeating units of PPS. A sample ^1^H-NMR spectrum is displayed in Supporting Figure 4A. ^1^H-NMR (400 MHz, CDCl_3_) δ: 3.6 (s, 68H, PEG), 3.3 (s, 3H, PEG-OCH_3_, reference), 2.9 (m, 2H/unit = 160/2 = 80H, -S-CH_2_CH(CH_3_)-S-), 2.6 (m, 1H/unit = 80H, -S-CH_2_CH(CH_3_)-S-). Additionally, GPC verified the conjugation of DP by demonstrating an increase in the polymer’s molecular weight (Supporting Figure 4B,C).

### Electrophoretic mobility shift assay (EMSA)

The loading of mRNA into PPDP2 nanocomplexes was evaluated using EMSA.^21,22^ EMSA was used to determine when ∼100% mRNA was loaded, and this ratio was used for all subsequent experiments. For sample preparation, 1 μg of mRNA was complexed with 40 μg of PPDP2. A 10 μL aliquot of these nanocomplexes was mixed with a loading buffer and then loaded onto a 1% agarose gel that contained GelRed® nucleic acid stain and was submerged in Tris-acetate-EDTA (TAE) buffer (40 mM Tris-base, 20 mM acetic acid, and 1 mM sodium EDTA). Electrophoresis was carried out at a constant voltage of 100 V for 30 minutes (Bio-Rad, Inc.), and the gels were subsequently imaged using a LAS 4010 Gel Imaging System (GE Healthcare).

### Cryogenic transmission electron microscopy (cryo-TEM)

Before plunge-freezing, 200-mesh copper grids with a lacey carbon membrane (EMS Cat# LC200-CU-100) were glow discharged using a Pelco easiGlow (Ted Pella) at 15 mA for 30 s under 0.24 mbar pressure. This process imparted a negative charge on the carbon membrane to promote even sample distribution. Next, 4 µL of a 5 mg/mL sample (either PPDP2 or PPDP2 + mRNA) was applied to the treated grid, blotted for 5 s with a +1 blot offset, and then plunged into liquid ethane using an FEI Vitrobot Mark IV. The grids were stored in liquid nitrogen. For imaging, the grids were loaded into a Gatan 626.6 cryo transfer holder and examined at –175 °C in a JEOL JEM1400 LaB6 emission TEM at 120 kV using a Gatan OneView 4k camera.

### Small Angle X-ray Scattering (SAXS)

SAXS measurements were conducted at the 5-ID beamline of the DuPont–Northwestern–Dow Collaborative Access Team (DND-CAT) at the Advanced Photon Source (APS) of Argonne National Laboratory. Throughout the experiment, each sample was exposed to collimated X-rays at a wavelength of 1.24 Å (9 keV). The samples, prepared at a 2 mg/mL concentration, were introduced into an in-vacuum flow cell and held in quartz capillaries of 1.6 mm thickness. Scattering profiles were collected in the q-range of 0.0015–0.08 Å ¹, with a sample-to-detector distance of approximately 8.5 m and an exposure time of 5 s. Calibration of the beamline was performed using silver behenate and a gold-coated silicon grating (7200 lines/mm). The momentum transfer, q, was defined by q = (4π/λ)sinθ, where 2θ is the scattering angle. Subsequent data reduction and buffer subtraction were executed with BioXTAS RAW,^23^ and model fitting was performed using SasView 5.0.5.

### Nanocarrier Formulation and characterization

The resulting sterile PPDP2 was mixed with *rrsp*-mRNA, *egfp*-mRNA, or CleanCap® *mCherry*-mRNA (TriLink BioTechnologies, San Diego, CA) in 25 mM sodium acetate buffer at the weight ratio of 40:1 to assemble monodisperse spherical complexes for mRNA delivery or for tracing transfection through the expression of fluorescently-labeled proteins after successful transfection, respectively. For quality control, PPDP2-*egfp*-mRNA, PPDP2-*rrsp*-mRNA, and PPDP2-*mCherry*-mRNA nanocarriers were characterized using a Nano ZS Zetasizer (Malvern Pananalytical), Azure 600 (Azure Biosystems), cryo-TEM (JEOL JEM1400), and SAXS (Advanced Photon Source) to ensure consistent nanocarrier diameter & charge (Figure 7A,B), nucleic acid loading efficiency (Figure 7C), morphology, and structure. For Zetasizer and SAXS measurements, 5% dextrose was used as the solvent to prevent salt interference with PPDP2.^24^ To validate the size and morphology, cryo-TEM studies were performed. The cryo-TEM images confirmed that PPDP2 self-assembled into uniformly spherical nanostructures and that upon mRNA incorporation, the nanoparticles retained their spherical morphology while exhibiting an increased diameter, consistent with DLS measurements (Figure 7D,E). SAXS analyses, modeled using a core–shell spherical fit in SasView software, further revealed that mRNA incorporation significantly expanded the predicted particle diameter from 24.9 nm to 61.1 nm and that both PPDP2 and PPDP2 + mRNA conformed to the core–shell spherical model, confirming their spherical nanostructures (Figure 7F). Sample data of validation are shown in Figure 7.

**Figure 7.**
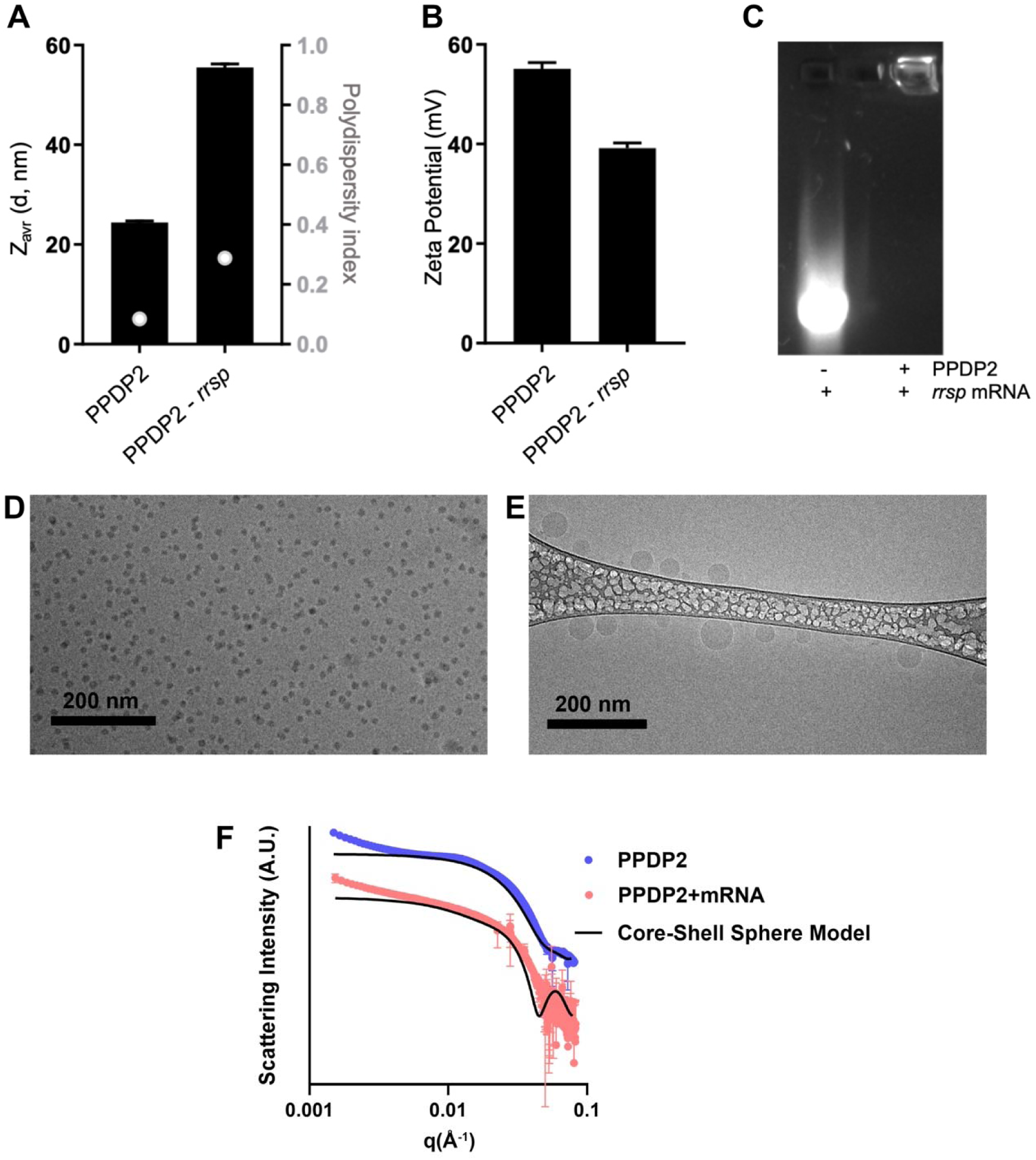
Sample data on PPDP2 loading of *rrsp*-mRNA. (A) Z-average size of PPDP2 alone and PPDP2 loaded with *rrsp*-mRNA. Polydispersity index (*y*-2 axis, grey) of PPDP2 alone and PPDP2 loaded with *rrsp*-mRNA shown as white/grey marker. (B) Zeta potential of PPDP2 alone and PPDP2 loaded with *rrsp*-mRNA. (C) Gel electrophoresis of naked *rrsp*-mRNA and PPDP2/*rrsp*-mRNA nanocomplexes at a 40:1 wt. ratio, demonstrating retention and immobility of mRNA within wells following stable complexation with PPDP2. (D,E) Representative cryo-TEM micrographs of D) PPDP2 and E) PPDP2 + mRNA, Scale bar = 200 nm. F) SAXS analyses on both PPDP2 and PPDP2 + mRNA, modeling was performed with a core–shell spherical fit in SasView.

### Immunofluorescence

PANC-1 cells were plated on 4-well slides in DMEM (10% FBS, 1% P/S) media and allowed to attach overnight. The following day, 1 µg *egfp*-mRNA was mixed with 40 µg of PPDP2 in ddH_2_0 for 30 minutes at room temperature or 1.5 µL MessengerMax Lipofectamine (Invitrogen) in media without FBS as a positive control according to manufacturer’s instructions. The mixture was added to cells along with negative controls, *egfp*-mRNA and nanocarriers alone. After 24-hour incubation, cells were washed three times for 5 minutes with phosphate-buffered saline (PBS), fixed with 100% methanol for 10 minutes, washed three times for 5 minutes with PBS, and mounted with 4′,6-diamidino-2-phenylindole (DAPI) stain. Cells were visualized on a Nikon Ti2 widefield microscope using a 40x objective. Five images were taken for each treatment and the total number of cells were counted per frame and compared to the number of cells that were expressing EGFP.

### SDS–PAGE and Western blotting

After treatment for 3 hours with the indicated concentrations of *rrsp*-mRNA or *rrsp*\*-mRNA by PPDP2 nanocarriers, fresh media was exchanged, and cells were harvested 21 hours later (24 hours total treatment). Protein extracts were prepared by either directly adding 2X sodium dodecyl sulfide (SDS)-polyacrylamide gel electrophoresis (PAGE) buffer to the tissue culture well or by harvesting cells by adding radioimmunoprecipitation assay (RIPA) buffer (50 mM Tris-HCl, pH 7.4, 150 mM NaCl, 2 mM Ethylenediaminetetraacetic acid (EDTA), 1% Nonidet P-40, 1% sodium deoxycholate, 0.1% SDS, supplemented with cOmplete mini protease inhibitor cocktail (Roche, catalog. no 11836170001) and 1 mM phenylmethylsulfonyl fluoride (PMSF). Equal amounts of proteins or equal volumes were separated by SDS-PAGE followed by Western blot analysis as described previously.^25^ Membranes were blotted using the following antibodies: anti-pan-RAS (Thermo Fisher Scientific, catalog no. MA1-012, RRID:AB_2536664), which recognizes RAS Switch I and thus detects only uncleaved RAS including mutant forms of RAS. Anti-GAPDH (Cell Signaling Technology, catalog no. 2118S) (as indicated) was used for normalization. Secondary antibodies used were fluorescent-labeled IRDye 680RD goat anti-mouse (LI-COR Biosciences, catalog no. 926-68070, RRID:AB_10956588) and IRDye 800CW goat anti-rabbit (LI-COR Biosciences, catalog no. 926-32211, RRID:AB_621843), respectively. Blot images were acquired using the Odyssey Infrared Imaging System (LI-COR Biosciences) and quantified by densitometry using NIH ImageJ software (ImageJ, RRID:SCR_003070). Percentage of uncleaved RAS was calculated as described previously.^9^

Protein extracts from frozen tissues were prepared by pulverizing tissue with mortar and pestle and homogenizing tissue in a microcentrifuge tube containing RIPA buffer. Samples were homogenized on ice three times, 5 seconds each time, incubated on ice for 30 minutes, and centrifuged at 22,400 x*g* for 15 minutes at 4 °C. Supernatants were collected and protein content measured using the BCA protein assay kit (Thermo Fisher Scientific) according to manufacturer’s instructions.

### Crystal violet assays

Cytotoxicity was assessed by staining cells with crystal violet. Briefly, 2 x 10^4^ cells/well were cultured in 24-well plates and treated with either 0.25, 0.5, 1, 1.5 or 2 µg *rrsp*-mRNA by PPDP2 nanocarriers or 2 µg *rrsp*-mRNA via MessengerMAX lipofectamine (Invitrogen) for 72 h. Cells were washed and crystal violet fixing/staining solution was added for 20 minutes at room temperature as described previously.^9^ Images of air-dried plates were acquired using a conventional desktop scanner.

### In vivo tumors

Mouse studies were conducted with female *nu/nu* mice at 6-8 weeks of age (Jackson Laboratories, Bar Harbor, ME) under protocols approved by the Northwestern University Institutional Animal Care and Use Committee. Cell line-derived xenograft tumors were initiated in the Northwestern University Developmental Therapeutics Core Facility (DTC RRID:SCR_017948) by subcutaneous injection of 2 x 10^6^ cultured PANC-1 PDAC cells to the dorsal flank of five mice for each group. Sample size was determined based on power analysis of expected 20% growth of control compared to 30% reduction in treatment group. When tumors reached an average size of 80-120 mm^3^, mice were first randomized into four groups each with five mice to equalize the range and average initial tumor sizes and body weight in each group, the treatment option was then assigned randomly to each group, and i.t. treatment was initiated. Treatments included PPDP2 synthetic nanocarriers alone in 25 mM sodium acetate buffer, 0.25 mg/kg of PPDP2 + *rrsp*-mRNA, the third 0.25 mg/kg of PPDP2 + *rrsp*\*-mRNA, or PPDP2 + *mCherry*-mRNA. Both tumor size and mouse body weight were measured just prior to new injections. Pooled data for two independent studies with *rrsp*-mRNA are reported at Day 22. Experiment 1 was continued until day 27, when mice were euthanized, tumors excised, and either snap frozen in liquid N_2_ for western blotting or fixed in 10% formalin overnight for IHC. Experiment #2 was continued to day 55 (without treatment) when tumors were measured and mice were euthanized. The investigators were not blinded to the group when measurements were made.

### Histology, IHC, and image analysis

Paraffin-embedding, sectioning, hematoxylin and eosin (H&E) and IHC staining of mouse xenograft tissue specimens were performed by the Robert H. Lurie Comprehensive Cancer Center (RHLCCC) Mouse Histology and Pathology Core Facility. Tumor sections (5 µm) were used for H&E staining or IHC staining with anti-cytokeratin 19 [(CK-19), #ab76539; Abcam], anti-Ki-67 (#GA626; Dako), anti-pan-RAS [(RAS), #PA5-85947; Thermo Fisher Scientific], anti-Phospho-p44/42 MAPK [(ERK1/2; Thr202/Tyr204, (D13.14.4E) XP, #4370; Cell Signaling Technology], anti-mCherry (Rockland Immunochemicals, catalog. no. 600-401-P16) antibodies, which are described previously.^9^ Primary antibodies were detected using the appropriate secondary antibodies and 3,3’-diaminobenzidine revelation (Agilent Dako). For IHC analysis, ImageJ was used to count positive cells. Original images were converted into three separate channels. Stained channel was selected and RGB thresholds were adjusted to select only stained cells, removing the background. Selected area was analyzed and quantified as maximum intensity minus minimum intensity. Five images from each group were analyzed and averaged to indicate % positive cells.

### Statistical analysis

Graphpad Prism v.10 software was used for statistical analysis. Bar plots represent the mean of at least three independent experiments and the standard deviation (SD) or standard error of the mean (SEM) as indicated in figure legends. Statistical significance was assessed as described in figure legends.

### Author Contributions

The manuscript was written through contributions of all authors. All authors have given approval to the final version of the manuscript.

### Funding Sources

This work was funded by Chicago Biomedical Consortium Accelerator Award (to K.J.F.S) and funding from the SQI Synthesizer Research Grant Program (to E.S.). Core services were provided by the Northwestern University Mouse Histology and Phenotyping Laboratory, the Center for Developmental Therapeutics, the Advanced Microscopy Facility, the Structural Biology Facility, and the Institute for Chemistry of Life Processes, which are supported by a National Cancer Institute grant to the Robert H. Lurie Comprehensive Cancer Center (NCI P30 CA060553). Additionally, we acknowledge the assistance of the BioCryo facility at Northwestern University’s NUANCE Center, which has received support from the SHyNE Resource (NSF ECCS-2025633), the International Institute for Nanotechnology, and Northwestern’s MRSEC program (NSF DMR-1720139). This work benefited from the use of the SasView application, originally developed under NSF award DMR-0520547. SasView contains code developed with funding from the European Union’s Horizon 2020 research and innovation program under the SINE2020 project, grant agreement No 654000. DND-CAT is supported by Northwestern University, The Dow Chemical Company, and DuPont de Nemours, Inc. This research used resources of the Advanced Photon Source, a U.S. Department of Energy (DOE) Office of Science User Facility operated for the DOE Office of Science by Argonne National Laboratory under Contract No. DE-AC02-06CH11357.

## ACKNOWLEDGMENT

Alfa Herrera is thanked for assistance with microscopy and Caleb Stubbs is thanked for contribution to the development of mRNA for use in experiments. We thank Eric W. Roth for cryo-TEM experiment.

## ABBREVIATIONS

PEG-PPS: poly(ethylene glycol)-b-poly(propylene sulfide)
PPDP2: PEG-PPS conjugated to a dendritic cationic peptide
RRSP: RAS/RAP1-specific endopeptidase
KRAS: Kirsten rat sarcoma
MARTX: multifunctional-autoprocessing repeats-in-toxin
EGFP: eukaryotic-optimized green fluorescent protein EGFP
IHC: immunohistochemistry
AF2: AlphaFold2
LNP: lipid nanoparticle

## Supporting Information

**Supporting Figure 1.**
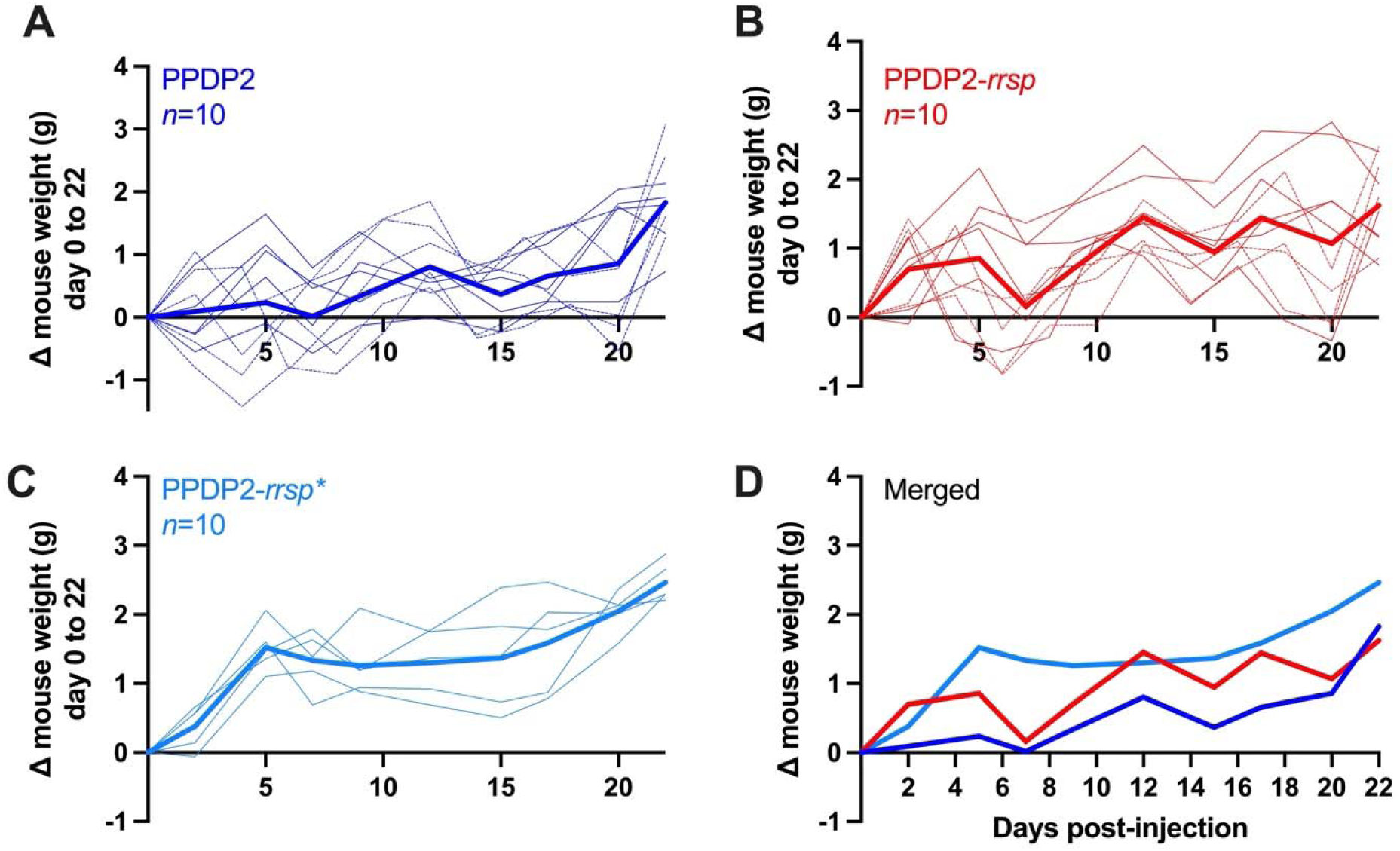
Change of weight of individual treated mice over time. (A-C) Mouse weight across days 0-22 for all individual mice inoculated as indicated. Solid thin line from experiment #1 and dashed lines from experiment #2. Heavy line is mean for merged experiments with points limited to days when mice were weighed in both experiments. (D) Merge of average line point, duplicate of Figure 4F.

**Supporting Figure 2.**
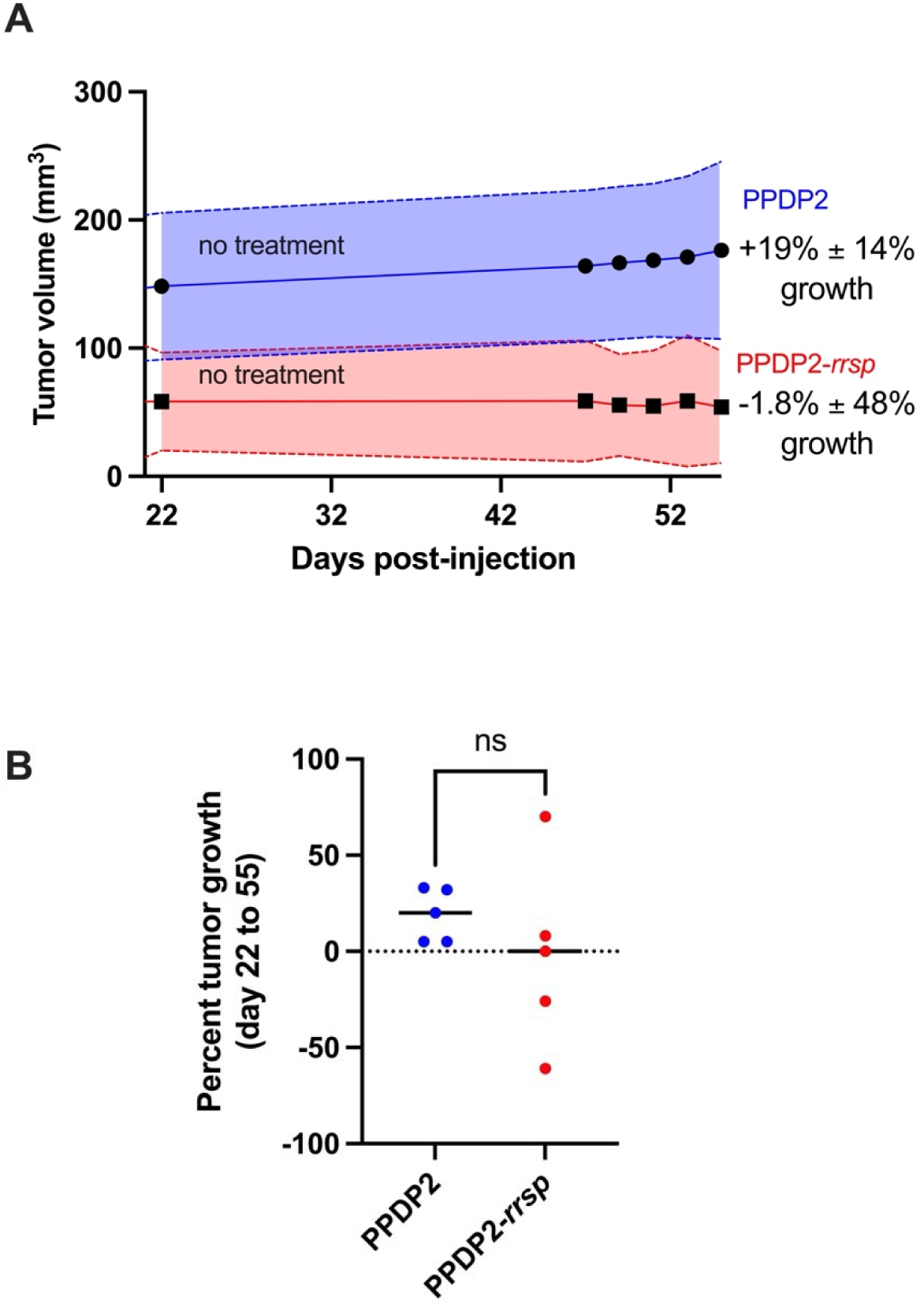
PANC-1 xenograft tumor volume for Experiment 2 extended without treatment from day 22 to day 55. (A) Mean +/- standard deviation of tumor volume from day 22 to day 55. Percent variation shown at right. (B) Percentage tumor volume change from day 22 to day 55. Data are not significantly different (as determined by Student’s *t* test) due to large variation of results in treatment group with range of change from +70% and +8% (growth) to -61% and -26% (regression). One tumor did not change (marked at 0) as it had fully resolved by day 22 and did not regrow.

**Supporting Figure 3.**
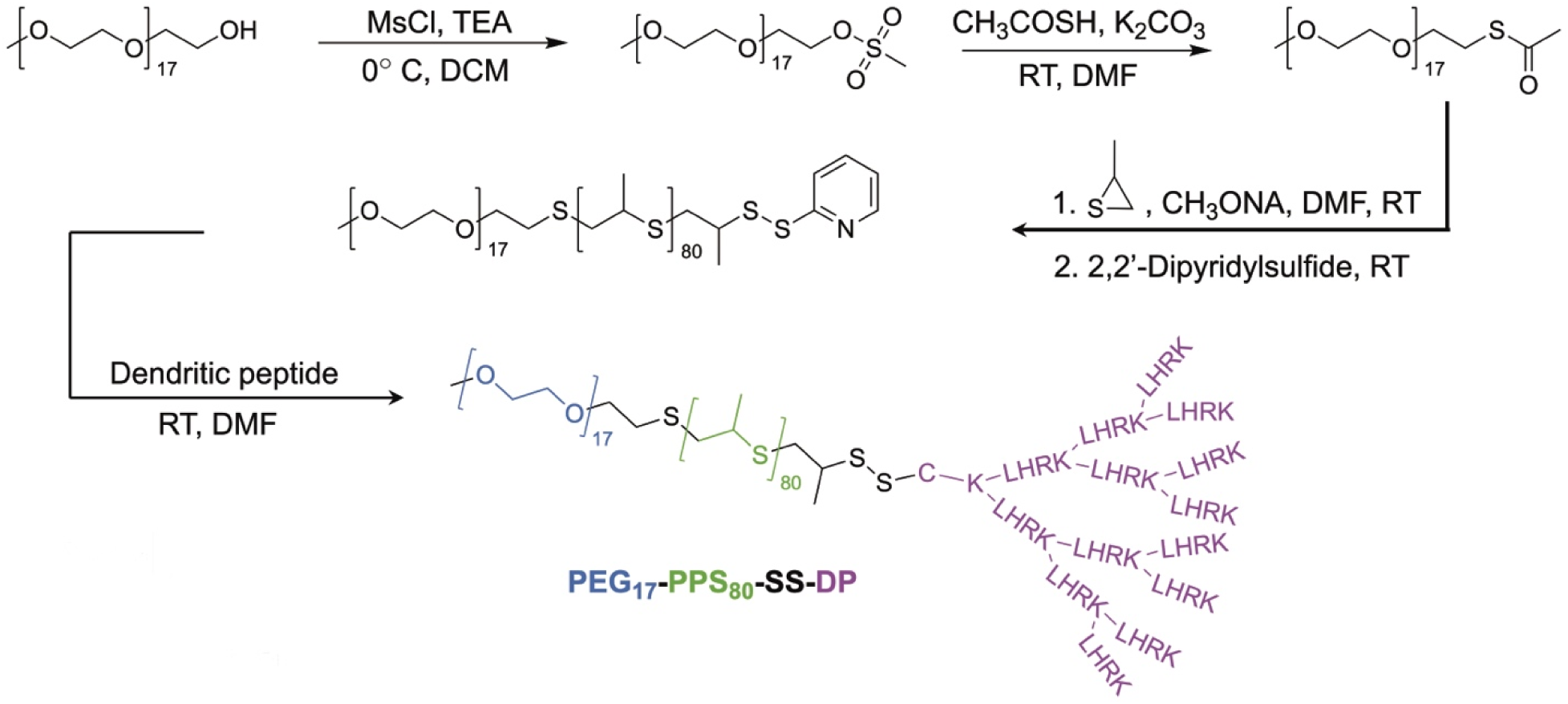
Reaction scheme for PPDP2. A schematic illustration outlining the synthesis process of the PEG_17_-b-PPS_80_-ss-Dendritic Peptide (PPDP2) polymer.

**Supporting Figure 4.**
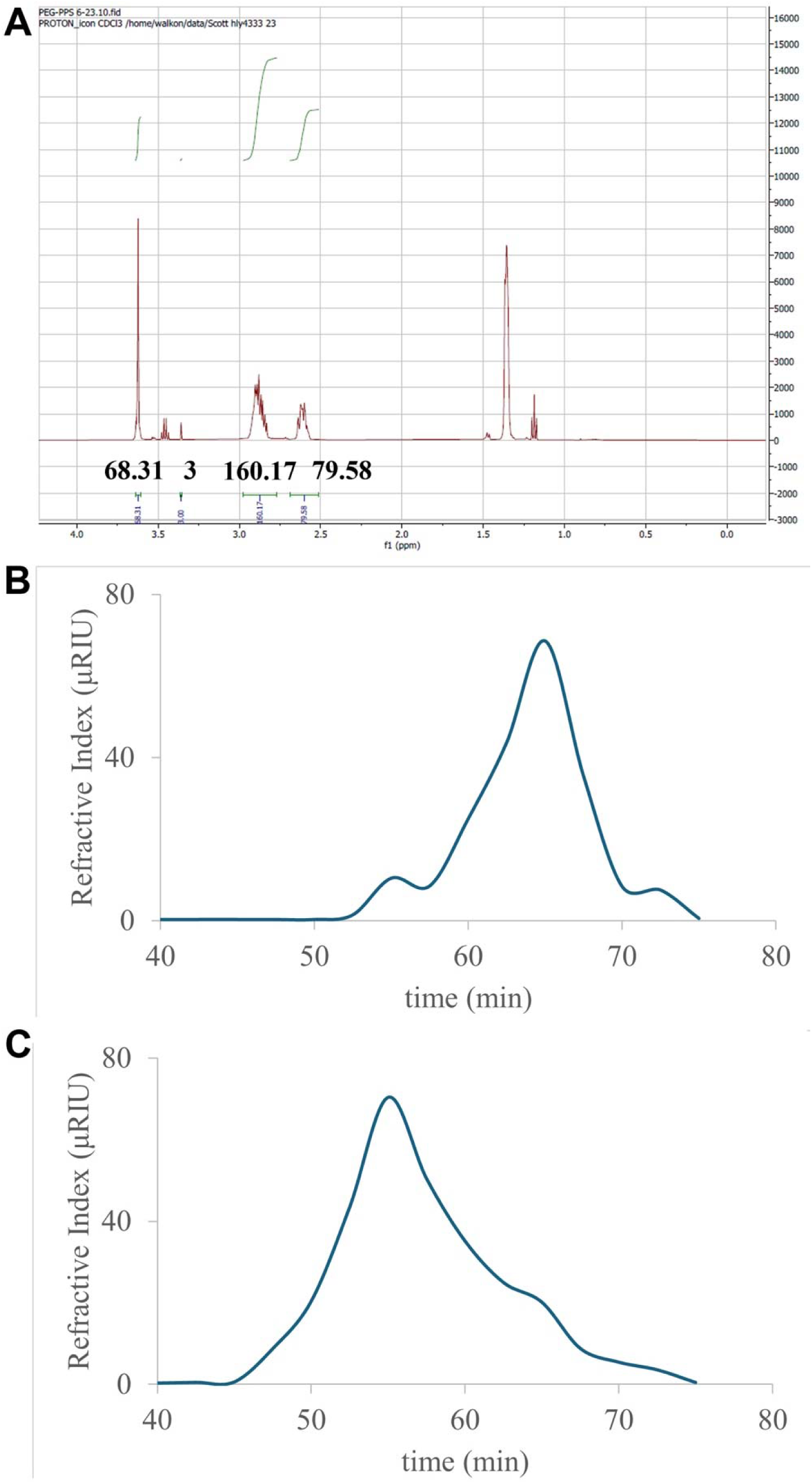
^1^H-NMR and GPC analysis of PPDP2. (A) ^1^H-NMR (400MHz, CDCl_3_) δ: 3.6 (s, 68H, PEG), 3.3 (s, 3H, PEG-OCH_3_, reference), 2.9 (m, 2H/unit = 160/2 = 80H, -S-CH_2_CH(CH_3_)-S-), 2.6 (m, 1H/unit = 80H, -S-CH_2_CH(CH_3_)-S-). (B,C) GPC performed for (B) PEG_17_-b-PPS_80_-pds and (C) PEG_17_-b-PPS_80_-ss-DP (PPDP2) was analyzed using PLgel columns equipped with refractive index and UV-Vis detectors, with a mobile phase composed of 10 mM lithium bromide in *N,N-*dimethylformamide.

**Supporting Table 1.**
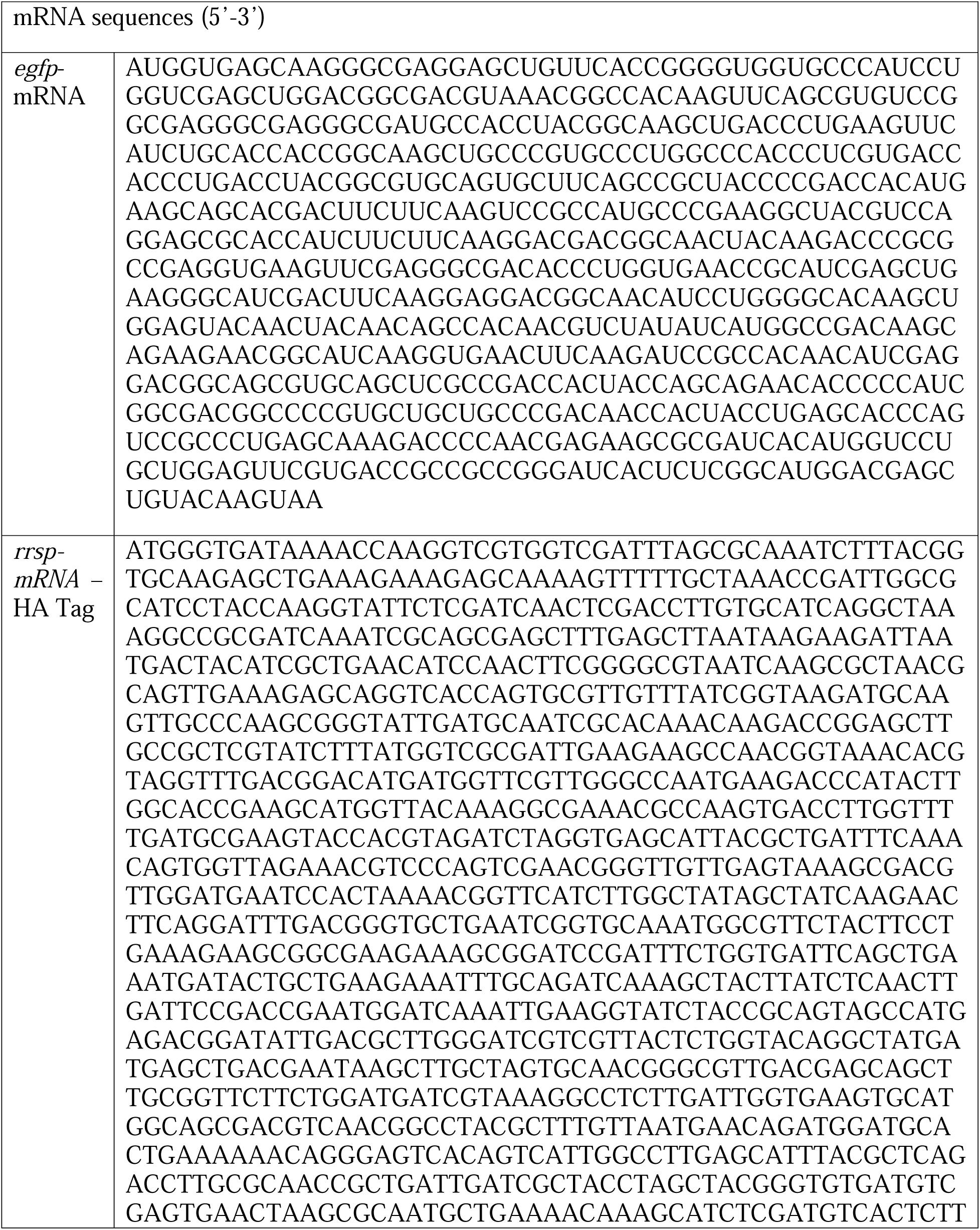
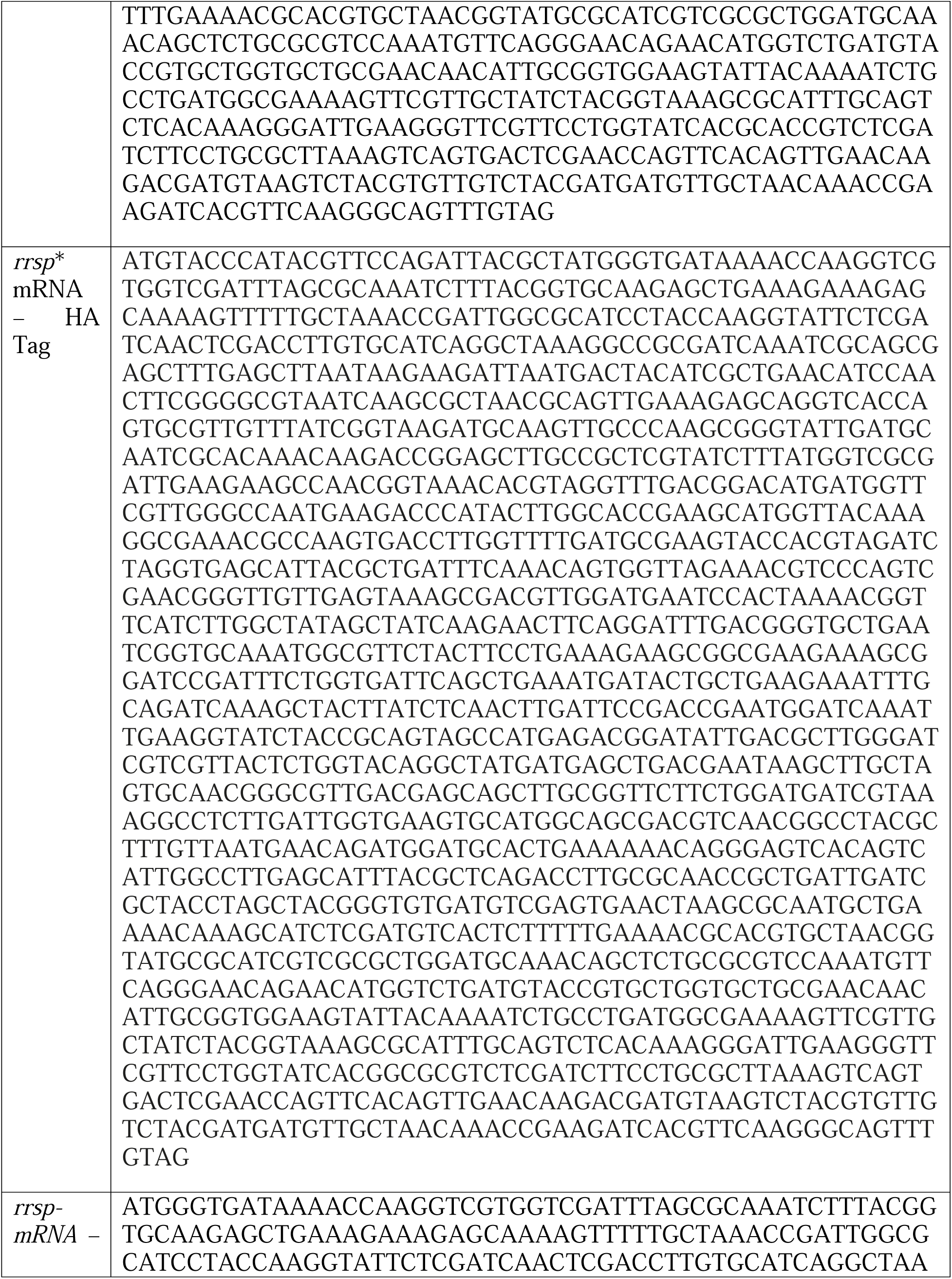
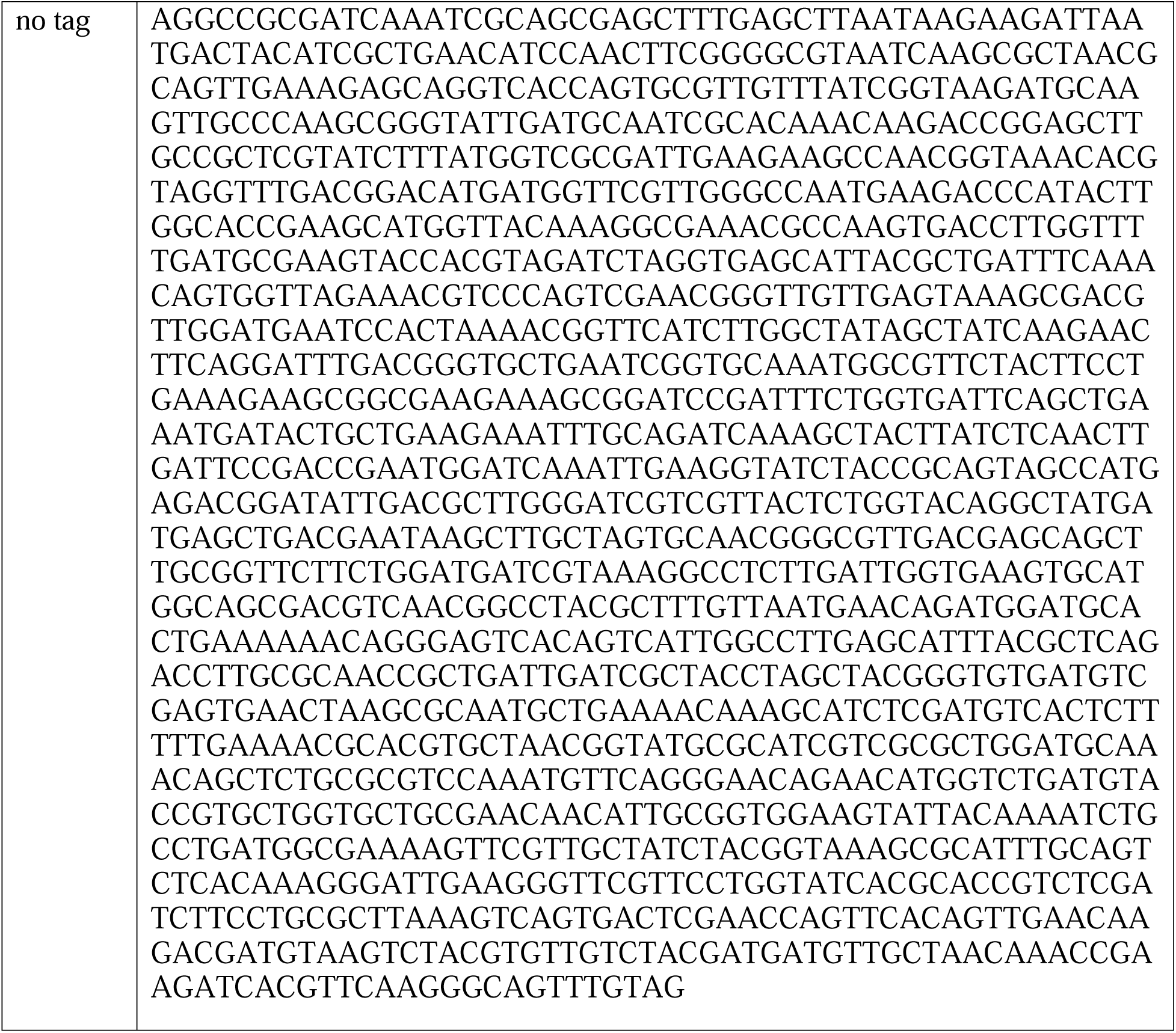
Sequences of mRNA molecules commercially synthesized for this study.

